# The universal labeling and sorting of avian primordial germ cell utilizing *Lycopersicon Esculentum* lectin

**DOI:** 10.1101/2024.05.09.593277

**Authors:** Hiroko Iikawa, Mizuki Morita, Aika Nishina, Yuji Atsuta, Yoshiki Hayashi, Daisuke Saito

**Affiliations:** Graduate School of Systems Life Sciences, Kyushu University; Department of Biology, Faculty of Science, Kyushu University, Fukuoka, Fukuoka, 819-0395, Japan

**Author notes:** Correspondence (D.S.).

## Abstract

Avian species serve as vital resources in human society, with the preservation and utilization of these species heavily reliant on primordial germ cells (PGCs). However, methods for effectively isolating these rare cells alive from embryos remain elusive in avian species beyond chickens. Even within chicken species, existing techniques have shown limited efficiency. In our study, we present a rapid, simple, and cost-effective method for labeling and sorting circulating-stage PGCs across various avian species, including Carinatae and Ratitae, utilizing *Lycopersicon Esculentum* lectin. Significantly, this method demonstrates high sorting efficiency by identifying a wide range of PGC subtypes while preserving the proliferative potential of chicken PGCs. This approach is anticipated to make a significant contribution to the conservation, research, and agricultural industries related to birds worldwide.

## INTRODUCTION

Avian species play crucial roles in various sectors including industry, agriculture, and scientific research. Consequently, conserving avian biological resources and implementing selective breeding and genetic manipulation are imperative. However, the established methods for cryopreservation of avian fertilized eggs, even in most common species like chickens, remain limited, and sperm cryopreservation techniques are also constrained (Svoradova et al., 2021). Although advancements are being made in avian embryonic stem cells (ESCs) and induced pluripotent stem cells (iPSCs) research (Dai et al., 2014; Pain et al., 1996), challenges persist in utilizing these undifferentiated cells for lineage preservation and the generation of genetically modified birds.

In this context, primordial germ cells (PGCs), the precursors of reproductive cells, have emerged as valuable tools for germline preservation, selective breeding, and the generation of genetically modified birds. During early development, PGCs undergo long-distance migration to reach the gonads (Richardson and Lehmann, 2010). In avian species, PGCs utilize the circulatory vascular system as their migratory route (Saito et al., 2022; Swift, 1914). The circulating PGCs in birds offer the potential for generating germline chimeras through their collection from the bloodstream and their transplantation into recipient embryo’s vasculature (Saito et al., 2022; Tajima et al., 1993). These chimeras, harboring donor germ cells in their reproductive organs have been shown to transmit donor germ cells to subsequent generations (Tajima et al., 1993). Moreover, PGCs are amenable to cryopreservation (Moore et al., 2006). Recent advancements in chicken PGC cultivation techniques have facilitated generating genetically modified PGCs, thereby initiating the production of transgenic chickens through transplantation into embryonic vasculature (van de Lavoir et al., 2006; Whyte et al., 2015).

Despite the significant utility of PGCs, their limited abundance within embryos underscores the critical need for efficient techniques for their collection. However, several challenges must be addressed in collecting PGCs from avian species. For example, in chickens, a common approach involves labeling PGCs derived from bloodstream or gonads with SSEA-1 antibody, which recognizes specific glycans on the cell membrane of chicken PGCs (Jung et al., 2005), and subsequently isolating them by Fluorescence-activativated cell sorting (FACS) (Ichikawa et al., 2022; Mozdziak et al., 2005). This method has several drawbacks, including its time-consuming procedural complexity, which often leads to sample loss, and the inability to label all PGCs (De Melo Bernardo et al., 2012). Although isolation of PGCs would be more feasible and efficient in reporter transgenic chicken lines specifically labeling PGCs (Chen et al., 2023; Hagihara et al., 2020; Rengaraj et al., 2022), the global availability of such chicken lines is not guaranteed. Furthermore, in avian species other than chickens, the detection of PGCs using SSEA-1 is more limited. For instance, it has been reported that PGCs in the bloodstream and gonads of Carinatae birds including the Japanese quail, Mallard duck, Muscovy duck, and turkey, as well as in the bloodstream and gonads of bird such as Ostrich, cannot be stained with SSEA-1 (D’Costa and Petitte, 1999; Hassanzadeh et al., 2019; Jung et al., 2017). Therefore, compared to existing methods, the development of a more versatile, simple, rapid, efficient, universally accessible, and cost-effective technique would significantly enhance research progress. However, currently, no method meets all these criteria.

In this study, we have addressed these challenges with a comprehensive solution. Our approach involves the utilization of *Lycopersicon Esculentum* (tomato) lectin (LEL) for the labeling and isolation of avian PGCs. LEL, a protein capable of recognizing and binding to specific sugar chain, the poly-N-acetyl D-glucosamine sugar residues (Nachbar and Oppenheim, 1982), has been employed for this purpose. Our findings demonstrate that LEL selectively accumulates on the cell membrane of viable PGCs across a broad spectrum of avian species, ranging from Carinatae to Ratites. Notably, the precision of PGC recognition achieved with LEL is remarkably high, potentially exceeding that of DDX4 (one of the most common PGC markers). Furthermore, the labeling process is exceedingly rapid, with results obtained within 15 minutes, and involves minimal procedural steps, thereby mitigating potential adverse effects on PGCs and minimizing sample loss associated with traditional staining techniques. Moreover, our method exhibits high sorting efficiency. Critically, post-sorting, the germness and proliferation efficiency of chicken PGCs remain unaffected.

## RESULTS

### LEL specifically labels PGCs in E2.5 chicken blood samples

In our previous research, we successfully identified PGCs within Embryonic day 2.5 (E2.5) chicken embryos by injecting LEL into the embryo’s bloodstream (Saito et al.,2022). To compare the efficiency and specificity of PGC staining in the chicken blood at Hamburger and Hamilton’s stage (HH) 14-15 (E2.5) (Hamburger and Hamilton, 1951) between LEL and SSEA-1, we conducted co-staining of these two staining (Fig. 1A, B). Among the collected blood samples, we observed approximately 1.5 times more LEL positive cells than SSEA-1positive cells (55.0%; 330 SSEA-1 positive cells/600 LEL positive cells) (Fig. 1B). Notably, within the SSEA-1-positive cell population, there existed variability in SSEA-1 signal intensity (Fig. 1B), encompassing cells with weak signals that were also accounted for in the count. PGCs (Fig. 1F), characterized by their large size (approximately 14 μm in diameter) and distinctive features such as oil droplets and glycogen granules in the cytoplasm (Ando and Fujimoto, 1983; Macdonald et al., 2010), were identifiable through optical microscopy. While the majority of SSEA-1 positive cells displayed morphological features consistent with PGCs, some were clearly identifiable as blood cells, wherein the SSEA-1 signal was also evident in the cytoplasm (Fig. 1C). In contrast, the LEL-positive cell population exhibited uniform and strong LEL signal intensity, with all LEL positive cells exhibiting PGC morphology, and no binding observed in blood cells, indicating higher specificity for PGC recognition compared to SSEA-1. Additionally, all SSEA-1 positive cells exhibiting PGC morphology were positive for LEL. Given the reported heterogeneity in SSEA-1 antigen expression in chicken PGCs, indicating incomplete labeling (De Melo Bernardo et al., 2012), LEL staining not only surpasses SSEA-1 in PGC recognition but also suggests the ability to detect certain subsets of heterogeneous PGCs undetectable by SSEA-1.

**Figure 1.**
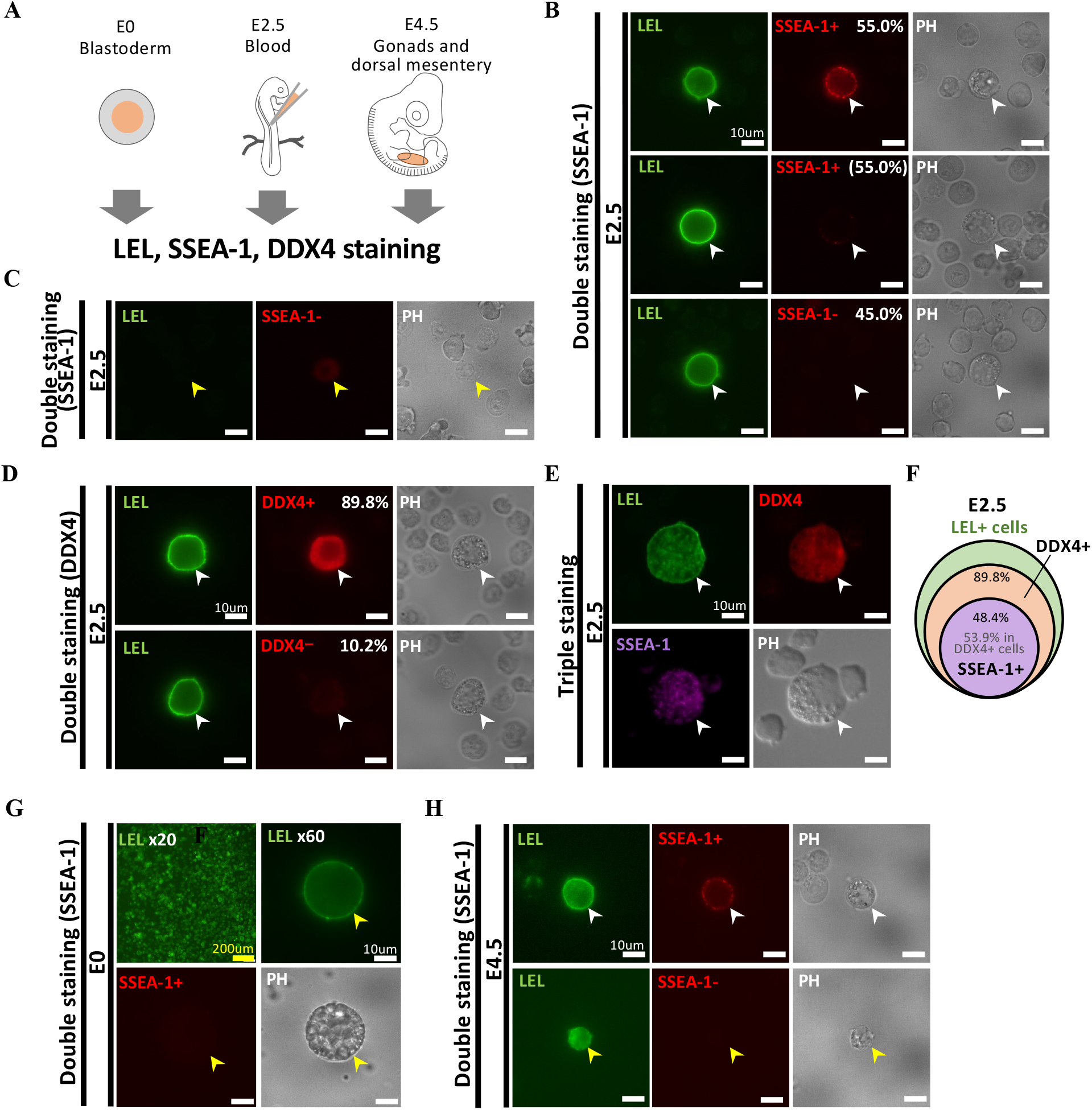
LEL specifically labels PGCs in E2.5 chicken blood samples. (**A**) The target tissues including PGCs are painted in orange at E0, E2.5, and E4.5, respectively. (**B-D**) Images of LEL staining with SSEA-1 or DDX4 signal in E2.5 chicken samples. The percentages represents the proportion of SSEA-1 positive (+)/negative (-) or DDX4+/- cells within the entire LEL+ cell population. (**E**) Images of triple staining with LEL+, SSEA-1+, and DDX4+ cells in E2.5 chicken samples. (**F**) The Venn diagram showing the relative abundance of SSEA-1+ and DDX4+ cells in LEL+ population. (**G, H**) Images of LEL staining with SSEA-1 signal in E0 and E4.5 chicken samples. White and yellow arrowheads indicate the cells presumed to be PGCs and other than PGCs.

To further validate the specificity of LEL binding to E2.5 chicken PGCs, we performed concurrent staining with DDX4, one of the most common PGC markers (Tsunekawa et al., 2000) (Fig. 1A). Blood samples collected from 20 embryos were stained with LEL, followed by fixation and SSEA-1 and DDX4 antibody staining (Fig. 1C, D). The number of LEL positive cells was approximately 10% higher than that of DDX4 positive cells (89.8%; 493 DDX4 positive cells/549 LEL positive cells) (Fig. 1D). Optical microscopic observation revealed that both DDX4 positive/LEL positive and DDX4-/LEL positive cells exhibited morphological features consistent with PGCs (Fig. 1D). The SSEA-1 positivity rate in DDX4 positive/LEL positive cells was 53.9% (55/102) (Fig. E, F). Given the reported heterogeneity in DDX4 expression in chicken PGCs (Rengaraj et al., 2022), LEL staining demonstrates the potential to detect subsets of heterogeneous PGCs undetectable by DDX4. Overall, LEL exhibits high specificity for E2.5 PGCs and suggests its capability to stain a significant portion of heterogeneous PGC populations.

In the subsequent investigation, we aimed to determine whether LEL could specifically label chicken PGCs at developmental stages other than E2.5. We conducted co-staining experiments using LEL and SSEA-1 antibody on samples at Eyal-Giladi and Kochav’s stage X (EGK stX; E0) and HH24-25 (E4.5) stages, representing pre- and post-PGC migration stages, respectively. In dissociated samples from E0 blastoderms, SSEA-1 staining was absent, suggesting that PGC recognition via SSEA-1 was not feasible at this stage. LEL signal was detected, with positivity observed in 70.7% of all examined cells (769 LEL positive cells/1088 cells, Fig. 1G). Considering the reported low number of PGCs at E0 (Tsunekawa et al., 2000), it is unlikely that LEL staining is PGC-specific at E0. In dissociated samples from E4.5 chicken gonads and dorsal mesentery of 15 embryos, LEL positive cells were 6.3 times more abundant than SSEA-1 positive cells. While most of SSEA-1 positive cells exhibited PGC morphology, only 11.6% (95 SSEA-1 positive cells/817 LEL positive cells) of LEL positive cells displayed PGC morphology (Fig. 1H). Therefore, in the gonadal and dorsal mesenteric regions of E4.5 chicken embryos, LEL also reacts with somatic cells, indicating low specificity in PGC recognition (Fig. 1H).

### LEL staining is effective for sorting chicken PGC without compromising their proliferation potential

To date, the sorting live chicken PGCs through fluorescence-activated cell sorting (FACS) has predominantly relied on techniques employing SSEA-1 staining (Ichikawa et al., 2022; Mozdziak et al., 2005), except in cases where PGC reporter transgenic chicken lines were utilized. However, as indicated by the results in Figure 1 and previous report (De Melo Bernardo et al., 2012), it’s evident that not all chicken PGCs can be efficiently stained using SSEA-1 (Fig. 1B, D, E). Given the quicker and simpler process of LEL staining with higher efficiency in PGC staining, it was anticipated to be a valuable tool for sorting circulating PGCs. Therefore, we sought to verify whether LEL staining could indeed be utilized for sorting PGCs by a cell sorter. We collected blood from 10 E2.5 chicken embryos, stained it with LEL, and attempted sorting using a cell sorter. Initially, we gated based on cell size and complexity to exclude most blood cells, followed by a second gate to sort cells with strong FITC fluorescence (Fig. 2A). We isolated a total of 370 LEL-positive cells from 10 samples (37 PGCs/embryo). Examination of sorted cells under bright-field microscopy confirmed that all cells exhibited the characteristics typical of large, lipid-rich cells, consistent with PGCs. Furthermore, immunostaining with DDX4 antibody on sorted cells revealed that 100% (27/27) of the cells were DDX4 positive, underscoring the highly effective nature of LEL staining for PGC sorting (Fig. 2B).

**Figure 2.**
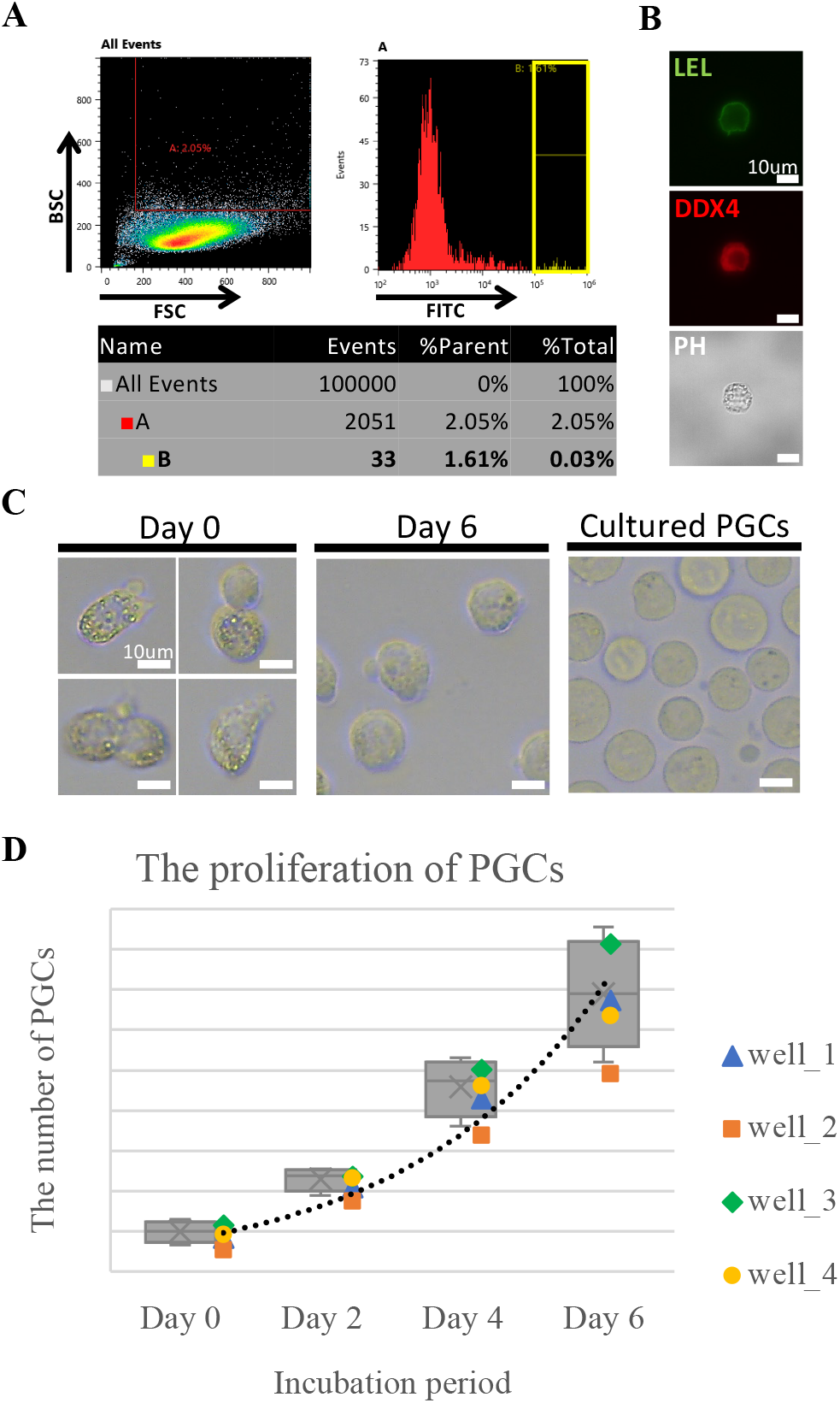
LEL staining is effective for sorting chicken PGC without compromising their integrity. (**A**) Dot-plots of forward scattering (FSC; x-axis) versus back scattering (BSC; y-axis), and histogram of FITC fluorescent. Chicken blood at E2.5 was sorted, and gate B (yellow square) was used for cell sorting. (**B**) Images of LEL staining with DDX4 immunostaining in sorted cells from E2.5 blood. (**C**) The images of culturing cells after cell sorting with LEL staining. Immediately following the cell sorting (Day 0) to 1 week after (Day 6), comparing to cultured PGCs (cultured for >100 days). (**D**) The box-plot of incubation period (day) versus the number of cultured PGCs after cell sorting. The dot line shows the approximate curve of these data.

Subsequently, we assessed the impact of LEL staining on the viability of chicken PGCs. Initially, PGCs sorted via LEL staining were cultured, and their proliferation and survival rates were evaluated. The initial week showed slow proliferation, followed by robust growth thereafter, mirroring the proliferation trend observed during culturing of PGCs derived from blood sample (Fig. 2C, D, Table1). This outcome indicates that LEL staining for PGC sorting does not impede PGC proliferation.

**Table 1.** The number of cultured PGCs after LEL+ sorting.

|  | Day0 | Day2 | Day4 | Day6 | Proliferation rate<br>(Day6 / Day0) |
| --- | --- | --- | --- | --- | --- |
| 1 | 19 | 45 | 91 | 142 | 7.473684 |
| 2 | 13 | 38 | 72 | 104 | 8 |
| 3 | 26 | 51 | 106 | 171 | 6.576923 |
| 4 | 21 | 50 | 98 | 134 | 6.380952 |

### LEL also specifically labels PGCs of quail, and emu at circulating stage

Reports of successfully isolating PGCs in avian species other than chickens have been scarce. This scarcity primarily arises from the inefficiency of SSEA-1 in identifying PGCs in non-chicken avian species and the lack of alternative techniques. Therefore, we sought to validate whether LEL could selectively recognize circulating PGCs in non-chicken avian species, specifically focusing on Japanese quail (Carinatae) and emus (Ratites).

Blood samples were obtained from embryos of these avian species at stages equivalent to HH14-15. Subsequently, LEL staining, fixation, and co-staining with SSEA-1 and DDX4 antibodies were conducted (Fig. 3A, B). In both samples, the presence of LEL-positive cells was consistently confirmed (Fig. 3A). Upon bright-field examination, these positive cells exhibited morphological features characteristic of PGCs, with no LEL signal detected in blood cells. When comparing SSEA-1 signal, in Japanese quail and emu samples, SSEA-1 signal was only observed in a small subset of LEL-positive cells (40.7% in quail, 38.5% in emu). Comparison of DDX4 and LEL signals revealed that DDX4 positive quail cells are included in a large subset of LEL positive cells (90.1%) similar to chicken PGC staining (Fig. 3B compared to Fig. 1C, D), and there are no DDX4 positive emu cells in LEL positive cells (0%) (Fig. 3B). This limited staining of DDX4 in emu PGCs may result from low cross-reactivity with antibodies generated against chicken DDX4, because N-terminal amino acids sequence, the epitope against this DDX4 antibody, is not conserved among species (Table2). Nonetheless, LEL exhibited high specificity in labeling circulating PGCs in Japanese quail and emus.

**Fig 3.**
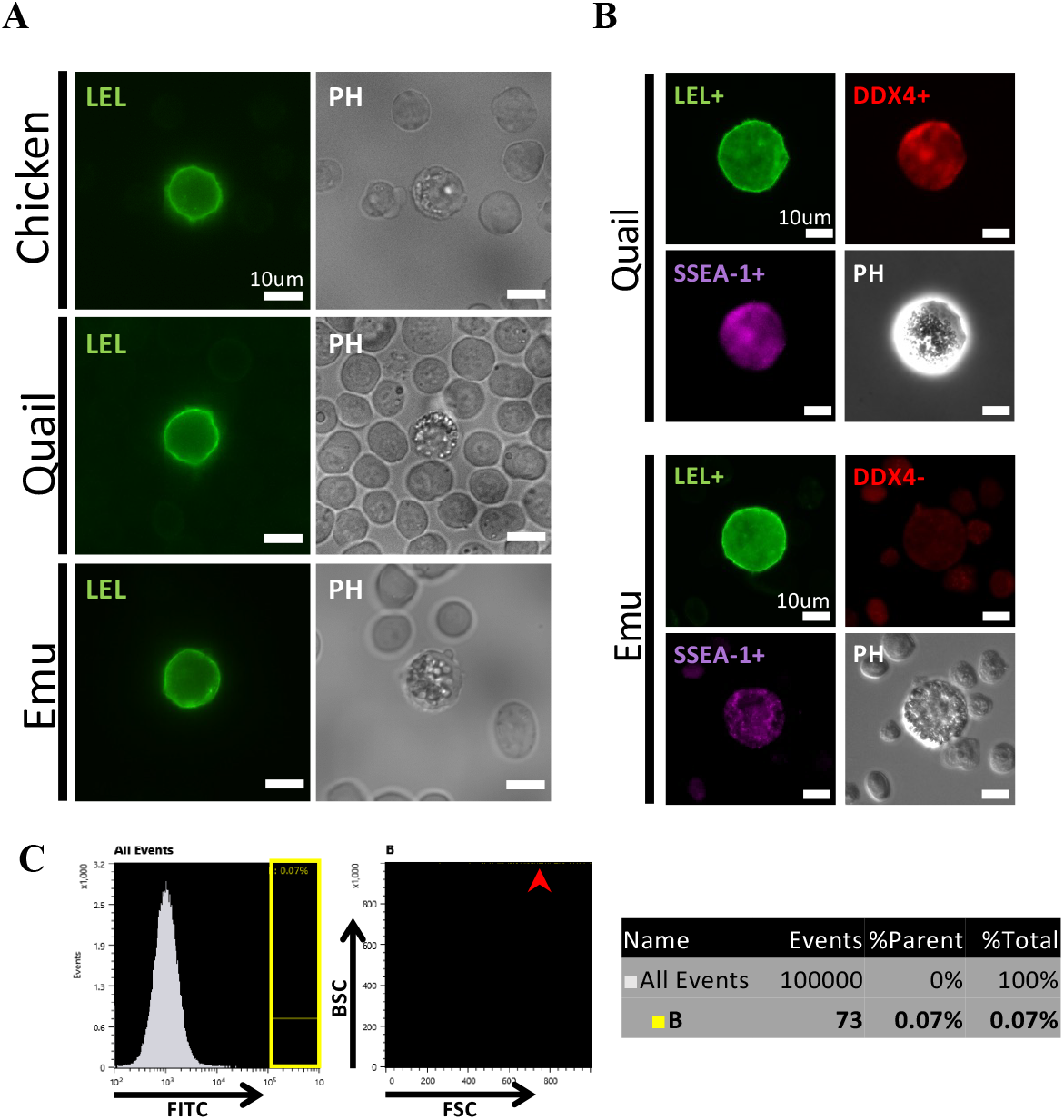
Blood PGCs from other avian species can be stained by LEL and purified by FACS. (**A**) The images of quail and emu blood PGCs stained by LEL. (**B**) Representative images of quail and emu PGCs, stained by LEL, DDX4, and SSEA-1. (**C**) Quail blood at HH14-15 was sorted and gate B (yellow square) was used for cell sorting. Histogram of FITC fluorescent and dot-plots of forward scattering (FSC; x-axis) versus back scattering (BSC; y-axis). Red arrowhead indicates the main subset of the distribution of the cells in gate B.

**Table 2.**
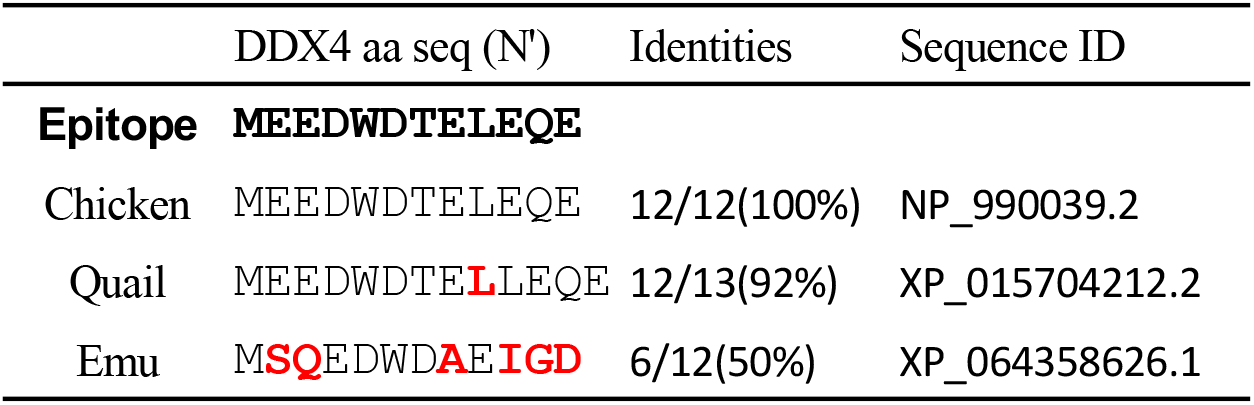
The differences of target sequences among avian species.

Finally, we explored the feasibility of sorting LEL positive PGCs using a cell sorter for Japanese quail. We collected blood from 10 HH14-15 quail embryos, stained it with LEL, and attempted sorting using a cell sorter. We gated to sort cells from LEL-stained blood from 10 HH14-15 quail embryos, with strong FITC fluorescence (Fig. 3C). We isolated a total of 115 LEL positive cells from 6 samples (19.2 PGCs/embryo). Examination of sorted cells under bright-field microscopy confirmed that all cells exhibited the typical morphology of PGCs (data not shown).

## MATERIALS AND METHODS

### Animals

Fertilized eggs from the BL-E strain of brawn leghorn chickens and the WE strain of Japanese quail were purchased from Nagoya University Avian Bioscience Research Center (Nagoya, Japan). Additionally, fertilized eggs were obtained from Yamagishi (Mie, Japan). Fertilized emu eggs were sourced from Kiyama farm (Saga, Japan). Chicken and Japanese quail eggs were incubated at 38.5°C, while emu eggs were incubated at 36.0°C in a high-humidity chamber (50-60%). All animal experiments were conducted in compliance with the ethical guidelines for animal experimentation established by Kyushu University.

### Sample collection and preparation for staining

Blastoderms, blood, and gonads (including dorsal mesentery tissue) were collected from EGK X, HH14-15, and HH24-25 embryos, respectively (Eyal-Giladi and Kochav, 1976; Hamburger and Hamilton, 1951). These samples were then placed in PBS on ice prior to the staining process. Blastoderms and gonadal tissues were minced and exposed to 0.05% trypsin/EDTA/PBS at 37.0°C for 10 minutes to facilitate dispersion after being washed once in PBS. During this dispersal process, cells were thoroughly dispersed by pipetting every 3 minutes. To ensure the retention of glycans on PGCs, the exposure time to trypsin should not exceed 10 minutes. The reaction was halted by adding an equal volume of FBS before centrifugation at 1500 rpm (approximately 138g). Prior to staining, the cells were washed and resuspended in PBS containing 0.2mg/ml of heparin (Sigma-Aldrich).

### LEL labeling

Before using, LEL DyLight® 488 (Vector Laboratories, Inc., DL-1174) was centrifuged at 15,000 rpm, 4°C for 30 minutes to remove the debris. In the following staining method, all the processes were conducted in shade on ice or at 4°C. Blood and dispersed blastoderm, gonadal tissues were incubated in 1/250 LEL in PBS for 5 minutes. To wash samples, 10 times quantity of cold PBS were added and centrifuged at 1500 rpm for 5 minutes twice.

### Co-staining with SSEA-1 and DDX4

In the case of SSEA-1 co-staining, cells were washed with PBS, and then incubated with a 1:100 dilution of anti-stage-specific embryonic antigen-1 (SSEA-1) with 5% FBS PBS for 1 hour in the shade on ice. After washing with PBS, the secondary antibody reaction was performed using donkey anti-mouse IgM and donkey anti-mouse IgG (Alexa fluor 568, invitrogen) at a 1:200 dilution with 5% FBS PBS for 30 min on ice. Subsequently, the cells were washed again with PBS and mounted on 96-well glass bottom dishes (AGC techno glass co., ltd) for observation. Photos were taken by sCMOS (ANDOR) attached to the macro zoom microscope IX83 (Olympus) with the software cellSens (Olympus).

In the case of DDX4 co-staining, cells were fixed with 4% paraformaldehyde in PBS for 30 minutes in the shade at room temperature after LEL staining. Following fixation, cells were washed twice with TNT (0.1M Tris, 0.15M NaCl, and 5ml Tween20), and then incubated in 1% blocking reagent TNT for 30 minutes in the shade. Subsequently, the cells were incubated with a 1:500 dilution of anti-DEAD-box helicase 4 rat IgG antibody (Atsuta et al., 2022) and a 1:100 dilution of anti-stage-specific embryonic antigen-1 (SSEA-1) mouse IgM antibody (sc-2170; Santa Cruz Biotechnology, Dallas, TX, USA) with 1% blocking reagent TNT for 1 hour on ice. After washing with TNT, the secondary antibody reaction was performed using donkey anti-rat IgG and donkey anti-mouse IgM (Alexa fluor 568 or 647, invitrogen) at a 1:200 dilution for 30 min on ice.

Subsequently, the cells were washed again and mounted on glass slides (Matsunami Glass Ind., Ltd) or 96-well glass bottom dishes (AGC techno glass co., ltd) for observation. Two color Photos were taken by sCMOS (ANDOR) attached to the macro zoom microscope IX83 (Olympus) with the software cellSens (Olympus). Three color photos were taken by ORCA-Flash4.0 (HAMAMATSU Photonics) attached to the macro zoom microscope DM6000B (Leica) with the software LAS X (Leica).

### Cell sorting

To isolate the LEL positive cells, SH800 (SONY) fluorescence activated cell sorter (FACS) was employed. Sorting parameters was automatically regulated at the biginning of the SH800 program. As an example of sorting parameters, the Droplet Clock was set to 30500 Hz, Droplet Drive to 19.01, Sort Delay to 30, Sort Phase to 0 degrees, Charge to 50.0, Deflection Left to -638, Deflection Right to 788. Chip alignment and calibration were conducted using the manufacturer’s fluorescent beads. Subsequently, cell sorter parameters were adjusted to utilize low FSC thresholds for detecting all particles in the sample (with a threshold of FSC set at 5.0%). Sensor Gain settings were adjusted to ensure accurate measurement of LEL staining (FSC = 4, BSC = 28% to 30%, and FL2 FITC = 30%). Throughout the sorting process, both the sample area and collection area were maintained at 4°C.

### Chicken PGC culture

The cultivation of chicken PGCs after cell sorting was carried out in FAcs medium in non-treated 96-well plates (Thermo, non-treated) (Chen et al., 2019; Saito et al., 2022). The FAcs medium composition includes 65.5 % DMEM (high Glucose, Ca-free, no glutamine) (Nakarai tesque), 21.8 % DDW(Fuji film), 1×Nucleotides (Millipore), 2.0 mM GlutaMax (Gibco), 1.2 mM Soldium Pyruvate (Gibco), 1×NEAA (Gibco), 55μM b-mercaptoethanol (Wako), 1% Penicillin-Streptomycin-Amphotericin (PSA) (Fuji film), 2% B27 Supplements (Gibco), 0.20 % Chicken serum (CS) (Biowest), 0.1 mg/mL Heparin (Sigma-Aldrich), 2 mg/mL Ovoalbmin (Sigma-Aldrich), 25 ng/mL ActivinA (API), 4 ng/mL FGF2 (Wako), 100 μM CaCl_2_ (Nakarai tesque). The PGCs were cultured at 38.0°C with 5% CO_2_ in a humidity incubator. Half of the medium was replaced with fresh medium every 2 days.

## DISCUSSION

In this study, we introduced a novel method for labeling avian circulating PGCs utilizing LEL staining. LEL swiftly binds to the cell surface of live PGCs through a simple procedure, displaying superior specificity in detecting a larger population of chicken PGCs compared to the conventional SSEA-1 labeling method. Importantly, our study revealed LEL’s capability to recognize PGCs across various avian species beyond chickens. The stained PGCs were effectively sorted using FACS, enabling precise PGC collection with remarkable accuracy. We demonstrated that the labeling and sorting process had no adverse effects on their viability, proliferation, and homing ability to the gonads of chicken PGCs.

### The heterogeneity of chicken PGCs and the recognition specificity of LEL

Previous single-cell RNA sequencing (scRNAseq) analyses of E2.5 Dazl::EGFP transgenic chicken PGCs revealed a distinct subset with low expression of DDX4 mRNA within the EGFP-positive cell population (Rengaraj et al., 2022). Furthermore, co-immunostaining of DDX4 and SSEA-1 unveiled a subset with reduced SSEA-1 staining among DDX4-positive cells (De Melo Bernardo et al., 2012). These findings highlight the heterogeneity in SSEA-1 antigen and DDX4 mRNA expression levels within the chicken PGC population, suggesting a decreasing number of PGCs expressing DAZL, DDX4, and SSEA-1 antigen in that order.

Our analysis revealed that all LEL-positive cells in the E2.5 chicken blood sample displayed morphological features consistent with PGCs and were more prevalent compared to DDX4- and SSEA-1-positive cells (Fig. 1). These findings indicate that LEL effectively identifies a substantial fraction of the heterogeneous PGC population. Notably, we achieved a chicken PGC sorting efficiency approximately 3.3 to 4.4 times higher (37 PGCs/sample) than that previously reported using SSEA-1 (8.45 to 11.1 cells/sample) (Ichikawa et al., 2022). This broad and robust recognition capability of LEL likely contributes to this enhanced PGC sorting efficiency.

Although accurately quantifying all PGCs in the sample remains challenges, it is reasonable to assume that LEL recognizes a broad spectrum of PGCs, including those expressing DDX4. Further comparative labeling analyses between DAZL and LEL may offer additional insights into PGC heterogeneity and further advance our understanding of PGC characterization.

### The universality of LEL binding to PGC surfaces

This study illustrates the potential of LEL for labeling and sorting PGCs in economically important avian species such as quail, and emu. Considering the lack of established techniques for specifically labeling live PGCs on the cell surface in these avian species, the successful cell sorting represents a notable advancement.

In samples obtained from HH14-15 quail and emus, every cell labeled by LEL exhibited PGC morphology. While the extent of LEL recognition towards heterogeneous PGC populations remains unclear in this analysis, it is apparent that LEL can label and sort a certain subset of PGC populations.

In all avian species investigated in this study, LEL effectively labeled PGCs equivalent to HH14-15 stages. This suggests that the glycans recognized by LEL on the PGC membrane surface may possess greater evolutionary conservation across species compared to those recognized by SSEA-1 antibodies. The preservation of LEL-recognized glycans in PGCs across diverse species is of significant interest. Considering their evolutionary origins in ancient ratites such as the emu as well as Carinatae birds, these glycans may also be conserved in neoaves such as the finches. Furthermore, if the presence of LEL-recognized glycans in PGCs is associated with their migration using the bloodstream, there is a possibility of their expression in the PGCs of certain reptiles that undergo similar movements (Golkar-Narenji et al., 2023). Hence, it is crucial to validate the suitability of PGC labeling using LEL across various animal species in future research.

